# Genome-Wide Association Studies meta-analysis uncovers *NOJO* and *SGS3* novel genes involved in *Arabidopsis thaliana* primary root development and plasticity

**DOI:** 10.1101/2023.07.17.549373

**Authors:** Brenda Anabel López-Ruíz, Berenice García-Ponce, María de la Paz Sánchez, Elena R. Álvarez-Buylla, Araxi O. Urrutia, Adriana Garay-Arroyo

**Affiliations:** Laboratorio de Genética Molecular, Desarrollo y Evolución de Plantas, Depto. de Ecología Funcional. Instituto de Ecología, Universidad Nacional Autónoma de México (UNAM); C. U. CDMX México.; Centro de Ciencias de la Complejidad, UNAM; Laboratorio de Genómica Evolutiva y Funcional, Instituto de Ecología, UNAM; Milner Centre for Evolution, Department of Life Sciences, University of Bath, Bath, BA2 7AY, UK

**Keywords:** *Arabidopsis thaliana*, accessions, primary root growth, GWAS, *SGS3*, *NOJO*

## Abstract

Postembryonic primary root growth relies on meristems that harbour multipotent stem cells that produce new cells that will duplicate and provide all the different root cell types. *Arabidopsis thaliana* primary root growth has become a model for evo-devo studies due to its simplicity and facility to record cell proliferation and differentiation. To identify new genetic components relevant to primary root growth, we used a Genome-Wide Association Studies (GWAS) meta-analysis approach using data published in the last decade. In this work, we performed intra and inter-studies analyses to discover new genetic components that could participate in primary root growth. We used 639 accessions from nine different studies and performed different GWAS tests ranging from single studies and pairwise analysis with high correlation associations, analyzing the same number of accessions in different studies to using the daily data of the root growth kinetic of the same research. We found that primary root growth changes were associated with 41 genomic loci, of which six (14.6%) have been previously described as inhibitors or promoters of primary root growth. The knockdown of genes associated with two of these loci: a gene that participates in Trans-acting siRNAs (tasiRNAs) processing *Suppressor of Gene Silencing* (*SGS3*) and a gene with a Sterile Alpha Motif (SAM) confirmed their participation as repressors of primary root growth. As none has been shown to participate in this developmental process before, our GWAS analysis identified new genes that participate in primary root growth. Overall, our findings provide novel insights into the genomic basis of root development and further demonstrate the usefulness of GWAS meta-analyses in non-human species.

## Introduction

*Arabidopsis thaliana* (hereafter Arabidopsis) is a plant distributed throughout Europe, Asia, and North America [1] and used as a model plant to study the molecular biology of natural variation because of its adaptation to local environments [2–4]. A multitude of Arabidopsis accessions have been collected around the world. many of which have been extensively characterised both phenotypically and genetically. In this sense, not only new genes have been discovered that participate in complex phenotypes such as primary root growth, but also it has been established the genetic interactions that underlie the evolution and development of phenotypic traits that enable plants to respond to the environment [2]. Due to the accessibility of an enormous set of sequenced accessions, there has been an increasing number of studies applying Genome-Wide Association Study (GWAS) strategies in this species[5]. GWAS has been used to study complex traits that result from a plethora of genetic variations of different populations that have evolved and adapted to diverse environments and help to identify the link between phenotype information and the genetic background [6]. Moreover, GWAS allows one to examine the statistical association between a particular phenotypic trait and a single nucleotide polymorphism (SNPs) [7].

Since the root is an organ necessary for proper anchorage, water and nutrients acquisition, microbial associations, and environmental perception [8], one of the phenotypic traits analyzed via GWAS is the primary root (PR) growth. The PR is highly plastic, and its length varies depending on the genotype and the environmental conditions where it grows[9]. There has been an increased effort to discover underlying genetic mechanisms involved in PR growth in response to soil contamination [10], nutrient availability changes, salinity stress [11,12] hormones response (13,14) and wound [15] using GWAS. In these studies, the SNPs associated with PR length under control conditions were not deeply analyzed because PR length is only used as a normalizer to study a particular stress condition or because small sample sizes render few or no statistically associated SNPs [11,14,16–18]. Nevertheless, the intense research on Arabidopsis PR has allowed uncovering some genetic mechanisms underlying root growth and differentiation as well as the response to different environmental conditions [19]. Several genes have been reported to influence PR, such as the transcription factors *WUSCHEL-RELATED HOMEOBOX 5* (*WOX5*), *PLETHORA* genes (*PLTs*), *SHORT ROOT* (*SHR*), *SCARECROW* (*SCR*), cell cycle genes members such as *CYCLIN-DEPENDENT KINASES* and *CYCLINS* and also plant hormones that are key regulators of cell division, elongation and cell identity, during PR development [19–21].

Here we reviewed different published studies and found that few of them focused their GWAS analysis on identifying genes that determine a differential PR growth between accessions under control conditions, that is, that does not involve exposure to a stress treatment. For instance, Slovak and collaborators (2020) found that the *ARABIDOPSIS ADENYLATE KINASE 6* (*AAK6*), a protein important for ribosomal biogenesis, is required for normal cell cycle progression and PR growth [22]. Furthermore, using an epistatic GWA analysis, Lachowiec and collaborators (2015) reported that the transcription factor of the NAC family *NAC6* and *ATL5, TERMINAL FLOWER1* (*TFL1*) and another locus that encodes an ankyrin family protein (At3g28880), contribute to PR length [23]. In addition, Justamante and collaborators (2019) reported five genomic regions that potentially participate in PR growth that contain a *CAMV MOVEMENT PROTEIN INTERACTING PROTEIN 7,* a gene that codifies a protein required for virus movement, a hypothetical gene (DUF810) that encodes for an unknown protein, *GRDP1* that is involved in germination and ABA response, a gene that encodes a hypothetical protein that participates in wound-induced lateral root development (*At4g01090*) and a gene that encodes a protein that participates in Arabidopsis senescence, *NON-YELLOW COLORING 1* (*NYC1*) [15]. Besides, we recently identified several genes that could regulate PR length as NAC domain-containing protein 48 (NAC048) and NAC3 [24]. Also, there are other studies for PR growth that found changes in the number and type of SNPs under salt stress using GWAS analysis [11,12].

In this study, we took advantage of the published data to try to use a huge number of accessions from studies to provide new genetic insights into PR growth. We performed different GWAS strategies to compare PR measures: a single GWAS using the data from all studies normalized by Z-score, and a case-control GWAS with 15% of the accessions with the highest or shortest PR length. We also carried out a pairwise analysis comparing the same number and kind of accessions in intra and inter-studies with high correlation. Interestingly we found a few accessions with the largest or the shortest primary root growth share between the nine different studies despite the lower correspondence between studies. Using all the different GWAS strategies, we found 41 loci that could participate in root development; of these, only 14.6% (6 genes) have been described as inhibitors or promoters of primary root growth opening new research possibilities. Moreover, we support our findings by analyzing two knockdowns of these loci: a gene that participates in Trans-acting siRNAs (tasiRNAs) processing *Suppressor of Gene Silencing* (*SGS3*) and a gene with a Sterile Alpha Motif (SAM) confirming their participation of both genes as repressors of primary root growth.

## Methods

### Data collection

We selected nine studies that evaluated the Arabidopsis PR length under control conditions (MS growth medium) [11–18,23] (S1 Table). The age of the seedlings was determined by calculating the number of days after sowing (DAS) based on the time the seeds were placed in the culture medium, as described in the methods section of each study. Also, we noted if the seedling age was indicated in days after germination (DAG). The data are represented as the mean PR length in cm (S2 Table) and the studies that had their data in pixels were transformed to cm according to [25] (1 pixel=21 μm) . The studies that displayed root kinetic growth as rate growth was converted to linear growth, adding to the previous value. From the PR length of each study, we calculated five different parameters, the mean of all accessions, the maximum and minimum values, the outliers date and the first and third quartile (S3 Table). The frequency of accessions used in each study is shown in S4 Table.

### Genome-Wide Association Studies

We checked the data for normality using the Shapiro-Wilk test, and the non-normal data were transformed by the Box-Cox transformation to resemble a normal distribution (S5 Table) [26]. The association between studies was tested by Pearson’s correlation coefficients using the PR length. The Genome-wide association mapping was performed using the GWAPP web interface, which contains genotypic information for ∼206,000 SNP markers [26] (http://gwas.gmi.oeaw.ac.at/). GWAS was conducted for PR length using the accelerated mixed-mode (AMM) to identify associations between the phenotype of the 639 unique accessions and the SNPs available in the database. Manhattan plots represent the genomic position of each SNP and its association [-log10(P-value)]. For each candidate gene, the annotations were retrieved from AraGWAS Catalog (https://aragwas.1001genomes.org/) and TAIR10 (Arabidopsis.org).

A single GWAS was performed with the mean of the 639 accessions data normalized by Z-score. The studies with root growth kinetics on average were carried out with the days evaluated. The jackknife method was used to estimate the bias of the Z-score average; for that, we recompute the GWAS leaving out one study at a time from the sample using only the average Z-score of the remaining data (S6 Table). Besides, we carried out GWAS using the media of the PR length for each study and selected the top 0.1% of 205,978 SNPs (n=205 SNPs for each study, 5,330 SNPs in total and searched for common SNPs (S7 Table).

Besides, 15% of the accessions with the highest or shortest PR length (S8 Table) were used in a case-control GWAS, where a binary system of accessions was assigned with the largest root a “1” and “0” for the smallest PR length. The GWAS analysis was performed through pairwise evaluation using the studies with high correlation (S9 Table), with the same number of accessions in different studies, or evaluating the root growth kinetic of the same research. Candidate SNPs were chosen by a minor allele frequency (MAF) ≥ 0.05. Genes located in the 10-kb window (5 kb up and downstream) of the associated SNPs were mapped and examined for natural variation.

### Polymorphism patterns in the selected genes in extreme accessions

Sequence data from the 1001 genome project [5] (http://signal.salk.edu/atg1001/3.0/gebrowser.php were used to analyze polymorphism among accessions with contrasting phenotypes using the 15% of accessions with short PR phenotype and the 15% accessions with long PR. SNPs located in the 10-kb window (5 kb up and downstream) of the associated SNPs were mapped and examined for natural variation (Horton et al., 2016). SNPs information of all available accessions was compiled and contrasted. To determine the effects of these variations and their exact positions, Variant Effect Predictor (VEP; [27]) was executed, with default parameters, on all SNPs found in these windows (https://plants.ensembl.org/Arabidopsis_thaliana/Tools/VEP. Those SNPs found in the largest number of significant accessions for each trait were counted to find SNPs with biological relevance.

### T-DNA insertion lines genotyping and phenotyping

The T-DNA insertion lines of the candidate genes were ordered from the European Arabidopsis Stock Centre (nasc.org.uk). The complete list of lines used is found in the Supplemental data (S10 Table). All mutant lines are in Col-0 background. The T-DNA insertion lines were genotyped by extracting DNA from leaf material ground in liquid nitrogen, an extraction buffer and isopropanol to precipitate the DNA and perform the PCR reactions. The primers used for T-DNA insertion line identification are listed in Supplemental Material (S10 Table).

For the analysis of the PR phenotypes of the T-DNA insertion lines, the seeds of Col-0 and T-DNA lines were stratified for 5 d at 4°C and grown vertically on Murashige and Skoog (MS) medium (0.2x MS salts, 0.05% MES, 1% sucrose, 1% agar, at pH 5.6) at 22°C under long-day conditions (16 h: 8 h, light: dark) with a mean light intensity of 976.121 ± 68.061 lumen/ft² measured with a HOBO device for 12 DAS.

### Quantitative reverse transcription PCR

RNA was extracted from Col-0, *sgs3-11, sgs3-13, nojo-1* and *nojo-2* complete roots of plants growing 12 DAS using Quick-RNA Miniprep (Zymo Research, Irvine, CA, USA). Complementary DNA was generated using the SuperScript^TM^II Reverse Transcriptase (Invitrogen^TM^) according to the manufacturer’s instructions. Real-time PCRs were performed using Maxima^TM^SYBR Green/ROX qPCR MasterMix (Thermo Scientific) on a StepOne ™ real-time PCR system (Waltham, MA, USA). The relative expression was calculated with three biological replicates controls with technical triplicates each one, using the 2^−ΔΔCt^ method, considering the Col-0 as reference and as internal housekeeping the *PROTODERMAL FACTOR 2* (*PDF2*; *AT1G13320*) and *UBIQUITIN-PROTEIN LIGASE 7* (*UPL7*, *AT3G53090*) [28]. All the primers used for these analyses are shown in Supporting Information S10 Table.

### Statistical analysis

The significance of SNP in GWAS was determined at the 5% FDR threshold computed by the Benjamini-Hochberg-Yekutieli method to correct for multiple testing [29]. Statistically significant difference in PR length was evaluated with ANOVA one-way followed by Tukey’s post-hoc tests. A value of p <0.05 was considered statistically significant (Graphpad Prism 8). For gene expression, statistical analysis was performed using the unpaired two-tailed Student’s t-test or one-way ANOVA followed by Tukey’s post-hoc tests, p <0.05 was considered statistically significant (Graphpad Prism 8).

## Results

### The experimental conditions determine the primary root growth plasticity

To discover novel genes involved in PR growth using the natural variation of Arabidopsis, we analyzed the PR length measures under control conditions of nine different studies to perform GWAS. Eight of these GWAS focus on identifying genes involved in PR growth under different stress conditions: salinity [11,12], nutrient deficiencies [16–18], hormones responses [13,14] and wound [15] and only one study is focused on obtaining SNPs under control conditions [23]. In some of these studies, the SNPs associated with the PR length under control conditions are not deeply analyzed may be due to a lack of statistical significance [11,13,14,16,18] or that the PR length was only used as a normalizer under a particular stress condition [17]. In two of these studies, significant SNPs were found under control conditions [12,15]; however, they employed the root growth rate as input for the GWAS or a linear model to perform the GWAS, making it difficult to compare the results with the ones obtained here. The number of accessions and the growth conditions used in these studies is shown in S1 Table, and the data for PR length is in S2 Table. Each study analyzed is represented with a capital letter and a number that indicates the days after sowing (DAS) or germination (DAG) used to measure the PR length. From the PR length of each study, we calculated five different parameters, the mean of all accessions, the maximum and minimum values, the outliers date and the first and third quartile (S3 Table) and the frequency of accessions used in each study for further analysis (S4 Table). We performed histograms representing the frequency distribution of the PR length of each study and checked the data for normality using the Shapiro-Wilk test and found that almost all the data have a normal distribution except the data of the day one evaluated by [16], the day 3 of [18] and all days evaluated by [12] (S5 Table, S1 Fig). Those data were further transformed into a normal distribution using the Box-Cox transformation to perform the GWAS according to [26].

Interestingly, we found that the mean PR growth reported in the nine studies is very different when comparing measures taken on the same day after sowing (DAS) time. For example, at 6 DAS (the capital letter indicates the study and the number of the DAS; see S1 Table), the mean PR length in cm was F6 (0.90 ± 0.16), H6 (1.04± 0.32) and I6 (0.46 ± 0.08) and so for 7 DAS (and other days; see S1 Table): A7 (4.46 ± 0.97), F7 (1.18±0,16) and I7 (1.10±0.19) (Fig 1A). These differences are statistically significant (ANOVA: p < 0.05, post hoc Tukey test: p < 0.05) except between F7 and I7. According to these, some PR lengths are similar when comparing different days of different studies; for example, C10 has a similar mean PR length with F13; D6 with I8 and D7 with F10 and so on (Fig 1A). The disparity in the mean primary root length could be because of the different accessions used, the sample size or/and the discrepancy in the growth conditions considering both the culture medium and the condition of the growth chamber. In these studies, as in many laboratories, Arabidopsis is usually grown on plates with gel-filled media. Still, each laboratory uses different concentrations of the components of the Murashige and Skoog (MS) medium and different agar concentrations and brands (S1 Table). It has been established that even in one laboratory using the same growth medium, the specific environmental conditions of the growth chamber are likely to vary to some degree affecting plant growth [30].

**Fig 1.**
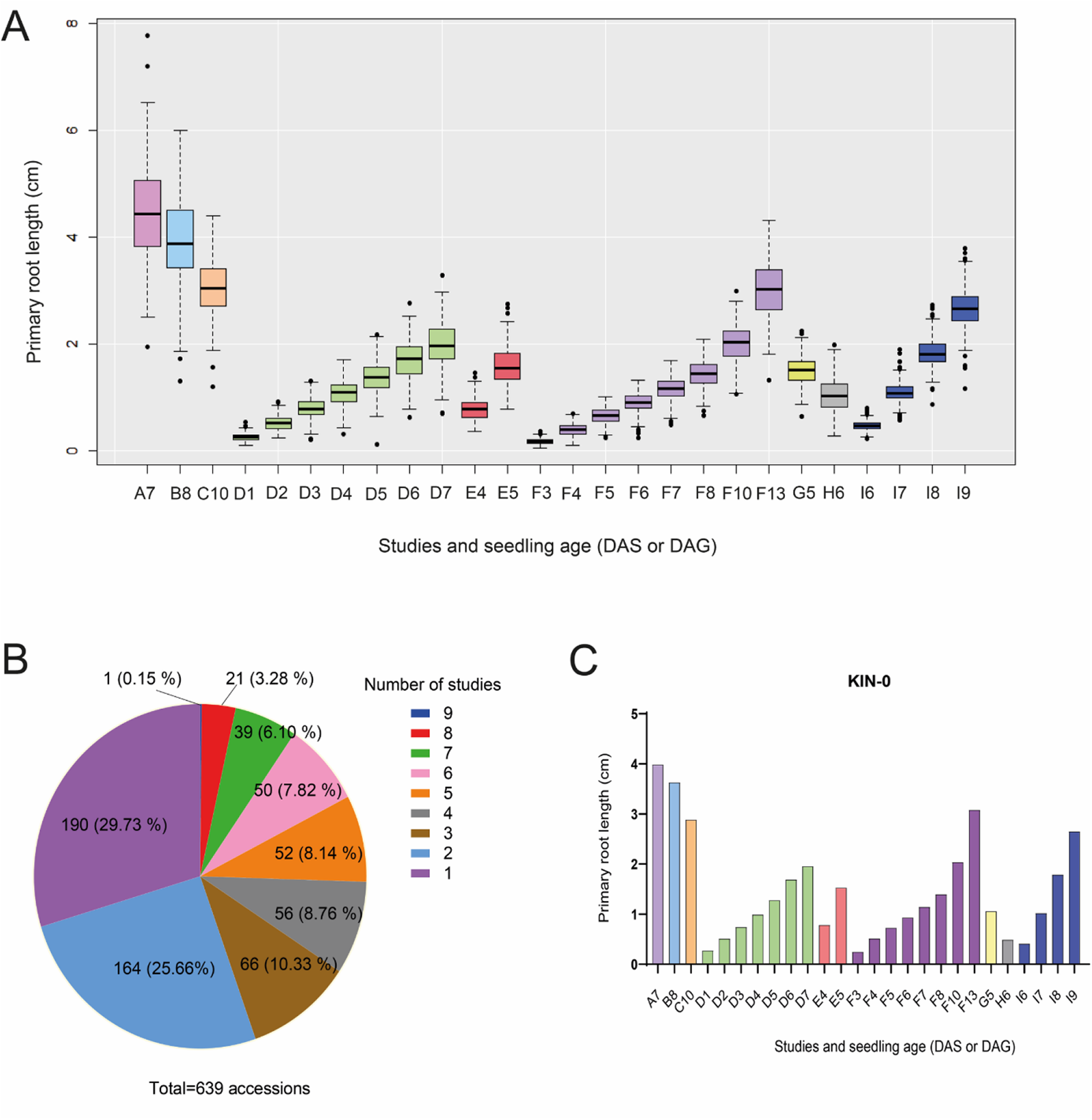
Comparison of the PR length between studies and the frequency of accessions used. (A). The box plot represents the distribution of the mean PR length of all accessions used in the different studies, the boxes indicate the first and third quartiles, the vertical line is the median, and the whiskers go from each quartile to the maximum or minimum value. The letter corresponds to each study, and the number means the DAS evaluated. Lachowiec et al., 2015 (A7) [23], Julkowska et al., 2017 (B8) [11], Ristova et al., 2018 (C10) [14], Li et al., 2019 (F3-F8, F10, F13) [18]; Justamante et al., 2019 (H6) [15], Deolu-Ajayi et al., 2019 (I6-I9) [12] or DAG: Bouain, et al., 2018 (D1-D7) [16], Bouain, et al., 2019 (E4-E5) [17], Ogura et al., 2019 (G5) [13], (B). Frequency of accessions shared between studies. (C). Primary root length of KIN-0, the only accession shared between all the studies and seedling ages.

When comparing all the studies, we found a correlation between the highest mean PR length, which was 4.46 cm (A7) from the study of Lachowiec et al. (2015) with the accession (VAR2-6) that has the largest PR in all the data we obtained (7.77 cm) [23]. Also, we found the same correlation between the lowest mean PR length F3 (0.17 cm) at 3 DAS from Li et al., (2019) with the shortest PR length of the STE-3 accession (0.051 cm; see S3 Table) [18]. In addition, STE-3 is also the shortest accession of other studies (D1, E4 and F3) despite the PR length of D1 (0.104 cm) being double the length of the PR of F3 (0.051 cm). Accordingly, with this disparity, the length of the shortest and largest PR varies greatly between different studies, even though similar seedling ages are evaluated. For example, CHR-1 is the accession with the shortest PR length in D5 (0.11 cm), D7 (0.68 cm) and G5 (0.64 cm). Moreover, TAMM-27 is the accession with the largest PR length in F4, F6 and H6. Nonetheless, the PR of H6 is larger (1.98 cm) than that of F6 (1.36 cm) (S3 Table). It is worth mentioning that, among the studies that have growth kinetics, the accessions that are the shorter or the larger change along the different days of the study [16,18]. For instance, in the work of Bouain et al. (2018) STE-3 is the accession with the shortest PR on day 1, COM-2 on day 2, and CHR-1 on days 3, 5 and 7, whereas on days 4 and 6 are HR-10 and CIT-0, respectively [16]. These results show the phenotypic variation of the root that responds dynamically to growing conditions.

Moreover, since PR length variation also depends on the sample size (number of accessions used), we analyzed the number of accessions shared between studies. Of 639 unique accessions, 190 (29.73%) are exclusive for one study, 164 (25.66%) are present in two studies, 66 (10.33%) in three investigations and from 50 to 56 accessions are shared between four to six studies (Fig 1B and S4 Table). We found 21 accessions commonly used in eight studies, and again, the PR length varies notably when we compare the same day (see S2 and S5 Tables). For example, the PR length of BOR-4 is very plastic as its growth is different in five studies: A7 (3.89 cm), I7 (0.83 cm), B8 (4.26 cm) and I8 (1.59 cm) (S2 Table). We also found that Kin-0 is the only accession shared between all nine studies (Fig 1C). Interestingly, its root length exemplifies the PR growth changes that rely on particular experimental growth conditions of the different studies, e.g. at 7 DAS, the root length shows a variation of up to 4 times among different experiments (compare Fig 1A and 1C).

In summary, different accessions and growth conditions can affect primary root growth. Accordingly, when the same accessions are evaluated at the same days between studies, they still display distinct PR lengths exhibiting the plasticity features of the PR.

### Different strategies to perform GWAS reveal different SNPs identified

To identify new SNPs with data from nine studies published, first we evaluated if our approaches could reproduce the same GWAS results in the published studies. To this end, we used the data from Bouain et al. (2018), and we found the same significant SNPs only on days 6 and 7 under control conditions [16] (Table 1).

In addition, we performed different GWAS strategies to obtain significant or common SNPs between the PR length of the different studies: i) a single GWAS using the data of all studies normalized by Z-score, ii) individual GWAS for all the studies, iii) a case-control GWAS with 15% of the accessions highest or shortest PR length, iv) pairwise analysis adjusting sample size and experimental conditions and v) making intra and inter analysis between the studies.

**Table 1.** List of Genes Identified by the different GWAS strategies in the 10 kb window (*significant statistically)

| Chr | SNP | Maximum log<sub>10</sub>(P) value | Gene(s) in a region around SNP (10 Kb window) | Type of analysis |
| --- | --- | --- | --- | --- |
| 1 | 29678615 | 5.29 | AT1G78930. Mitochondrial transcription termination factor family protein<br>AT1G78920 Vacuolar H<sup>+</sup>-pyrophosphatase 2<br>AT1G78922 Transmembrane protein<br>AT1G78940 Kinase with adenine nucleotide alpha hydrolases-like domain-containing protein | Z-score |
| 1 | 28381609 | 5.7 | AT1G75580. SAUR51 SMALL AUXIN UPREGULATED RNA 51<br>AT1G75590.1 SAUR52. | GWAS using 15% accessions with shorter PR length |
| 2 | 18106620 | 5.8 | AT2G43650. EMBRYO DEFECTIVE 2777<br>AT2G43670.1 Carbohydrate-binding X8 domain superfamily protein<br>AT2G43660. Carbohydrate-binding X8 domain superfamily protein<br>AT2G43680. IQ-domain 14 (IQD14); | GWAS using 15% accessions with larger PR length |
| 5 | 15776264 | 6.16 | AT5G39430 DUF1336 family protein<br>AT5G39420 CDC2C | Case-control GWAS |
| 4 | 1215616 | 7.23* | AT4G02735 F-box SKIP17-like protein<br>AT4G02733 F-box family protein<br>AT4G02740 F-box/RNI-like superfamily protein | Single GWAS (D6 and D7). Pairwise analysis (same accessions between D5 and D6) |
|  | 1215701 | 7.11* |  | Single GWAS (D6 and D7). Pairwise analysis (same accessions between D4 and E5, D3 and D, D4 and D5) |
| 5 | 8004085 | 6.39* | AT5G23720 PROPYZAMIDE-HYPERSENSITIVE 1 (PHS1)<br>AT5G23730 Transducin/WD40 repeat-like superfamily protein<br>AT5G23740 Ribosomal protein S11-beta | Single GWAS (D6 and D7) |
| 2 | 6550883 | 6.1 | AT2G15100 Gypsy-like retrotransposon<br>AT2G15110 hypothetical protein (Protein of unknown function, DUF601) | Pairwise analysis (same accessions between D1 and F4) |
| 1 | 11435245 | 5.20 | AT1G31860 Histidine biosynthesis 2<br>AT1G31850 S-adenosyl-L-methionine-<br>dependent methyltransferases superfamily<br>protein<br>AT1G31870 pre-mRNA-splicing factor of<br>RES complex protein | Pairwise analysis (same<br>accessions between D2 and<br>F5, D3 and F5, E5 and F5) |
| 4 | 1260826 | 5.44 | AT4G02820 Pentatricopeptide repeat (PPR)<br>superfamily protein<br>AT4G02810 FANTASTIC four-like protein<br>(DUF3049)<br>AT4G02830 hypothetical protein | Pairwise analysis (same<br>accessions between D3 and<br>G5, E5 and G5) |
| 3 | 18054594 | 6.06 | AT4G38620 MYB DOMAIN PROTEIN 4<br>AT4G38630 regulatory particle non-ATPase<br>10 | Pairwise analysis (high<br>correlation between D3 and<br>D4, D4 and D5) |
| 1 | 24811500 | 5.27 | AT1G66500 Pre-mRNA cleavage complex II<br>AT1G66490 F-box and associated interaction<br>domains-containing protein<br>AT1G66510 AAR2 protein family<br>AT1G66520 formyltransferase<br>AT1G66480 plastid movement impaired 2 | Pairwise analysis (high<br>correlation between D2 and<br>E4, D2 and D3) |
|  | 24810980 | 5.62 |  | Pairwise analysis (high<br>correlation between D3 and<br>D4) |
| 5 | 7947758 | 5.77 | AT5G23575 Transmembrane CLPTM1 family<br>protein<br>AT5G23570 SUPPRESSOR OF GENE<br>SILENCING 3 (SGS3)<br>AT5G23580 calmodulin-like domain protein<br>kinase 9 (CDPK9) | Pairwise analysis (high<br>correlation between D5 and<br>D6) |
| 1 | 5426133 | 6.05 | AT1G15760 Sterile alpha motif (SAM)<br>domain-containing protein<br>AT1G15757 Molecular chaperone<br>Hsp40/DnaJ family protein<br>AT1G15770 mediator of RNA polymerase II<br>transcription subunit<br>AT1G15772 mediator of RNA polymerase II<br>transcription subunit<br>AT1G15780 mediator of RNA polymerase II<br>transcription subunit 15a-like protein | Pairwise analysis (high<br>correlation between F6 and<br>F7, F7 and F8, F8 and F10) |
| 3 | 8573982 | 5.69 | AT3G23780 Nuclear RNA Polymerase D2A<br>(NRPD2A)<br>AT3G23790 AMP-dependent synthetase and<br>ligase family protein<br>AT3G23770 O-Glycosyl hydrolases family 17<br>protein | Pairwise analysis (high<br>correlation between F10_F13) |

### Single GWAS from data of all studies

In this analysis, we used the mean of root length of the 639 accessions normalized by Z-score. Due to the differences in PR growth owed to culture conditions between the studies (Fig 1A), the SNP (29678615) that we detected did not have a high score (5.29) and mapped inside the coding region (exon 1) of the AT1G78930 (*MTERF16*) gene that encodes a mitochondrial transcription termination factor family protein. Although, the Z-score normalization helps us to reduce the noise of inconsistent data due to their heterogeneous sources, the average Z-score could not reflect the effect of every single article. Therefore, to overcome this, we evaluated the contribution of each study to the final GWAS result through the resampling method Jackknife. To this end, we delete, at the time, the date of each study and perform GWAS with the average Z-score of the remaining data. Excluding the data of either Bouain et al. (2018), Julkowska et al. (2017) or Justamante et al. (2019), we obtained the same SNP 29678615 but with a higher score -log_10_(P): 5.94, 5.89 and 5.66 respectively (S6 Table). Contrary, if we did not consider the rest of the studies independently, the scores drop from 5.29 to 4.2 (see S6 Table for details), indicating that those studies contribute more to the final result of the mean Z-score. Since we did not obtain significant SNPs using all the data, our following approach was to perform individual GWAS for all the studies and compare them to look for common SNPs.

### Individual GWAS of all the studies display significant SNPs

We used the media of the PR length to carry out a GWAS for each study (S7 Table). We evaluated and selected the top 0.1% of 205,978 SNPs (n=205 SNPs for each study, 5,330 SNPs in total) and found 3,287 unique SNPs and 1,032 SNPs that are shared by two or more studies. Although we did not detect common SNPs among the nine studies, several SNPs are shared among them but with a low score (S7 Table). It is noteworthy that of the fourth SNPs found, one of them SNP 1215701 had the maximum frequency among studies; it is shared between three studies and their days evaluated: Bouain et al., (2018) (-log_10_(P) =7.11 (D7), 6.72 (D6), 5.99 (D5), 5.54 (D4), 4.60 (D3), 4.36 (D2), 3.46 (D1), Bouain et al., (2019) (-log_10_(P) = 4.32 (E4), 4.05 (E5)) and Ristova et al., (2018) (2.92) (S7 Table). he SNP 18016370 was found when comparing the study of Ristova et al. (2018) and Julkowska et al., (2017) ((-log_10_(P) =5.5 and 4.35 respectively; S7 Table). The SNP 16769779 is shared between D4-D7, E5, F7-F10, and G5, whereas 3559602 is a SNP shared between D2, D4-D7, C10, F7 and F10 (see S7 Table for scores).

### GWAS of the accessions with the shortest or largest primary root length

To execute a GWAS with contrasting groups of accessions, we select extreme accessions of each study, choosing the 15% of accessions with the shortest and largest PR length (S8 Table). For the studies that carried out root kinetics, we decided only to take the last day for the largest PR and the first day for the shortest PR. Besides, in the case of duplicate values, we select the shortest or largest value for the first or last DAG or DAS, respectively. For the GWAS with the largest PR length, the SNP with the highest score (-log_10_(P) = 5.8) was 18106620. This SNP is not found intra-genetically, and the nearest genes in a 10 kb window are the *EMBRYO DEFECTIVE 2777 (EMB2777)* gene, two genes that encode members of Carbohydrate-binding X8 domain superfamily protein and a gene encoding a protein IQ-domain 14 (Table 1). In addition, when using the shortest PR length data, we identify the SNP 28381609 with a -log_10_(P) = 5.7 and the two genes near this SNP, encoding for *SAUR51* and *SAUR52* (Table 1).

Since we eliminated the duplicate values between the studies and chose only the shortest or the largest PR length, and this could bias the final result, we decided to carry out a case-control GWAS selecting the accessions with the shortest and largest PR and giving them a value of 0 or 1 as we described below.

### Case-control GWAS

We implemented a case-control GWAS with the 15% PR largest and 15% lowest accessions of all the studies. For that purpose, we used a binary system of accessions, being “1” the largest and “0” the smallest in each study. 271 accessions were classified in this way, and the accessions shared between both groups were eliminated (S8 Table). With this approach, we found one SNP (15776264) with a -log_10_(P) = 6.16 score that maps in the coding region of the gene AT5G39430 that encodes a DUF1336 protein with an unknown function (Table 1). The low proportion of common SNPs found is probably owed to the number and the different accessions used (ranging from 93 to 347) (S1 Table), so we select the same accessions between studies to perform a GWAS to resolve these two issues.

### GWAS correction with sample size and experimental conditions

The sample size limits the power of GWAS since the phenotypic variation for a trait can overlap with the population structure leading to an artifact [7,31]. Our first approach was to select common accessions between studies to execute GWAS.

We performed pairwise analysis between studies that shared the same accessions and selected the top 0.1% SNPs. As can be seen in Table 2, we only compared studies where the number of shared accessions was higher than 100 and, in the case of studies with different days, we selected those days that had the highest correlation between studies. Of the five studies without kinetic data, only (B8) of Julkowska et al., 2017 and (C10) of Ristova et al., 2018 shared more than 100 accessions, we detected when comparing them, 8 SNPs of which we selected the one with the highest -log_10_(P) (3455993; Table 1). Furthermore, comparing D1 and F4, the highest -log(P) value (6.1) was obtained with the SNP (6550883) (Table 2). This SNP is mapped within the genomic region of a gypsy-like retrotransposon (AT2G15100). Interestingly, we found that in several pairwise analyses (D2 and F5, D3 and F5, E5 and F7), we obtained the same top SNP 11435245 that is positioned within the AT1G31860 Histidine biosynthesis 2 gene with a – log_10_(P) value ranging of 3.82 to 5.20 (Table 2). Likewise, we identified the SNP 1260826 that mapped in the gene AT4G02820 encoding a Pentatricopeptide repeat (PPR) superfamily protein when comparing D3 and G5 and E5 and G5. Otherwise, in the comparison between the F4 and I9 analysis, they do not have any SNP in common (Table 2).

**Table 2.** Pairwise GWAS between studies that shared the same accessions. The top 0.1% of common SNPs are indicated.

| Pairwise analysis | Number of accessions shared | Level of correlation | Number of SNPs shared | SNPs with the highest $-\log(P)$ | $-\log_{10}(P)$ in both studies | Gene associated |
| --- | --- | --- | --- | --- | --- | --- |
| **B8_C10** | 107 | 0.458 | 8 | 3455993 | 4.40, 5.77<br>4.40, 5.77 | AT5G10946 Hypothetical Protein<br>AT2G14750 Adenosine-5'-Phosphosulfate (APS) Kinase |
| **C10_E5** | 121 | 0.473 | 1 | 8004085 | 3.14, 3.66 | - |
| **C10_F3** | 108 | 0.1 | 4 | 19347877 | 3.43, 4.59 | AT3G52170 DNA Binding protein |
| **D1_F4** | 132 | 0.664 | 13 | 6550883 | 6.10, 4.13 | - |
| **D2_F5** | 132 | 0.727 | 25 | 11435245 | 4.70, 4.09 | AT1G31860 Histidine biosynthesis 2 |
| **D3_F5** | 132 | 0.725 | 26 | 11435245 | 3.82, 4.70 | AT1G31860 Histidine biosynthesis 2 |
| **D3_G5** | 129 | 0.421 | 9 | 1260826 | 3.98, 4.77 | AT4G02820 Pentatricopeptide repeat (PPR) superfamily protein |
| **E5_F7** | 132 | 0.75 | 16 | 11435245 | 3.84, 5.20 | AT1G31860 Histidine biosynthesis 2 |
| **E5_G5** | 128 | 0.437 | 4 | 1260826 | 4.36, 5.44 | AT4G02820 Pentatricopeptide repeat (PPR) superfamily protein |
| **F4_I9** | 109 | 0.159 | 0 | - | - | - |

### GWAS analysis using the same experimental conditions

As the developmental stage during which the phenotype is being studied is crucial to identify genes, we decided to perform a correlogram to check randomness in our data and to try to correct experimental conditions by choosing those values with high correlation and performing with them pairwise analysis intra and inter-studies (Fig 2). The relationship between each pair of studies was evaluated using Pearson correlation, where 0 is no correlation, 1 is a positive correlation, and −1 means a negative correlation. As expected, the highest association was found between the different days of the same studies: D1 and D2 (0.83), D2 and D3 (0.91), D3 and D4 (0.95), D4 and D5 (0.94), D5 and D6 (0.94), D6 and D7 (0.96), F3 and F4 (0.86), F4 and F5 (0.93) as the same for the next days and studies, except for I7 and I8 from Deolu-Ajayi et al. (2019) that correlate 0.76 (Fig 2). The subsequent high association was between the two separate studies of the same research group (Bouain et al., 2018; Bouain et al., 2019) (D and E; Fig 2). Interestingly, the results of Deolu-Ajayi et al. (2019) (I6-I9) show a negative correlation with the PR length reported by both Bouain et al., 2018 and Bouain et al., 2019 meaning that both articles are inversely related and the PR length cannot be compared between them. Likewise, I8-I9 have a low positive correlation with Julkowska et al., 2017 (B8), Ristova et al., 2018 (C10) and Li et al., 2019 (F3-F10). Thus, the data of Deolu-Ajayi et al. (2019) is the only study that does not have any positive association with the other studies (Fig 2) since it shows, on average, a faster growth rate and even when the same accession and days are compared. For instance, the accession KIN-0 has a growth rate of 0.61 cm/day between days 6 and 7, whereas for these same days in Bouain et al. (2018) and Li et al. (2019) the growth rate is 0.26 and 0.21 cm/day respectively.

**Fig 2.**
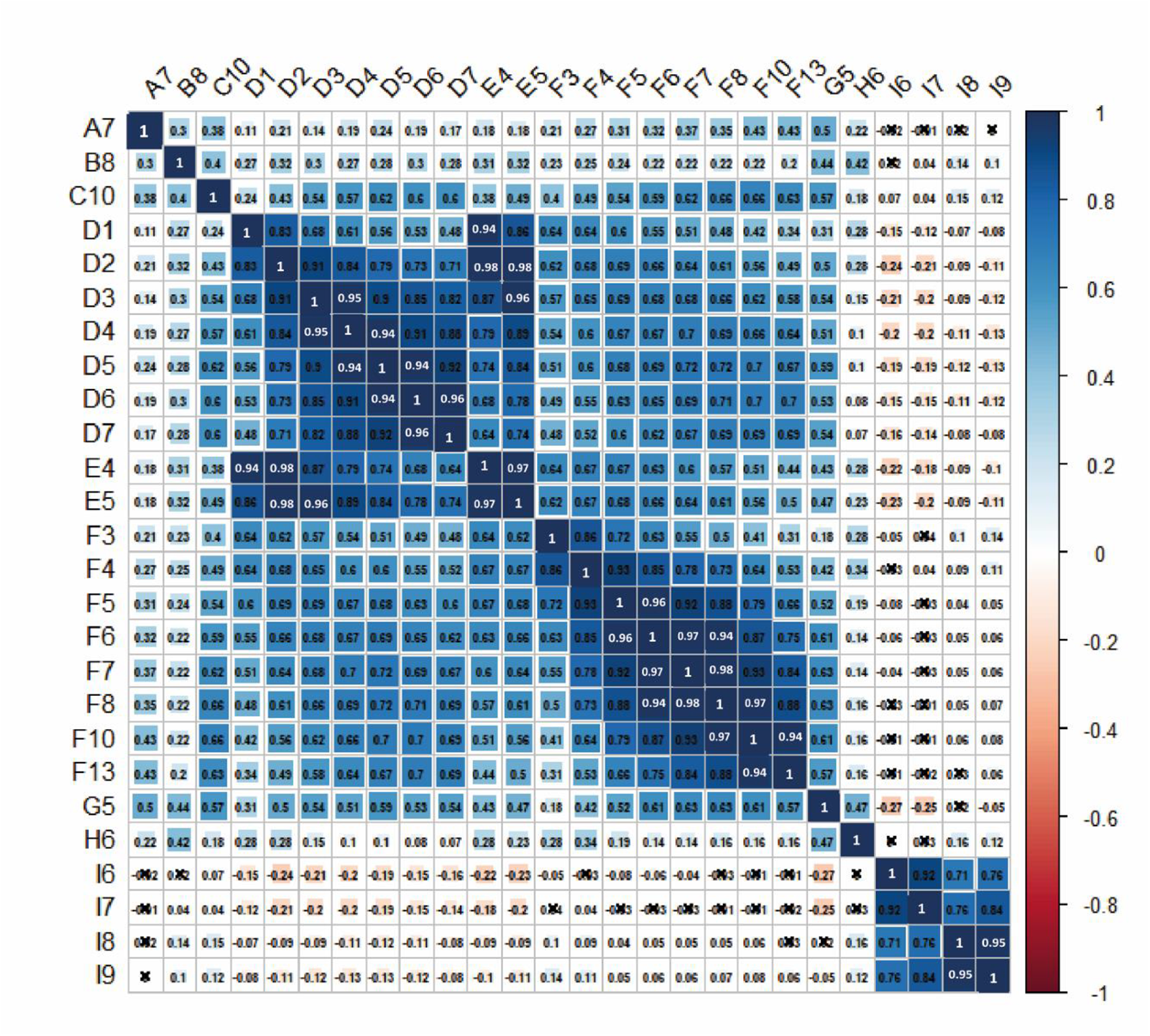
Correlation coefficients among the studies and the different seedling ages used in each one. The abbreviation of each study is depicted in S1 Table. On the right side of the correlogram, the legend colour shows the correlation coefficients and the corresponding colours; positive correlations are displayed in blue and negative correlations in red colour. The black cross in each cell depicts no association. The letters on the left side of the figure correspond to the different studies: Lachowiec et al., 2015 (A7); Julkowska et al., 2017 (B8); Ristova et al., 2018 (C10); Bouain et al., 2018 (D1-D7). Bouain, et al., 2019 (E3-E5), Li et al., 019 (F3-F8, F10, F13); Ogura et al., 2019 (G5), Justamante et al., 2019 (H6), Deolu-Ajayi et al., 2019 (I6-I9).

We selected those days with the highest correlation (0.9-1) to perform a pairwise GWAS analysis, and it was found that the proportion of common SNPs increased directly with the correlation (S9 Table) either intra or inter-studies. Furthermore, it was observed that in the root growth kinetics carried out by Bouain et al., 2018 (D2-D7), the SNPs shared rise as the seedling ages increment (S9 Table). With this approach and using the data from the study of Bouain et al. (2018) (D6 and D7), we identify four SNPs with the highest p-value, and that is above the threshold: 1215616, 1215701, 12662987 and 8004085 (-log_10_(P) =7.23, 7.11, 6.9 and 6.39 respectively (S9 Table and Fig 3A). The SNPs 1215616 and 1215701 are mapped in the coding region of the gene encoding AT4G02735 F-box SKIP17-like protein; 12662987 lies within a pseudogene, and 8004085 is localized in an intergenic region, and 3 genes were found within the 10-kb window of the mapped SNP (Table 1). In addition, the SNP 18054594 identified in the D3-E5, D4-E5 and in the D2-D3, D3-D4, D4-D5 pairwise studies, mapped in the intragenic region of *MYB DOMAIN PROTEIN 4* gene (Table 1 and S9).

Also, when we compared F6-F7, F7-F8, F8-F10, we discovered a SNP 5426133 (score 5.75) that mapped inside the exon of a gene that encoded to the Sterile Alpha Motif (SAM) domain-containing protein (Table 1 and S9). Another important SNP is 7947758 (score 5.7) which was obtained by analyzing the high correlation between D5 and D6. This SNP mapped to the 5th exon of the gene that encodes a transmembrane CLPTM1 family protein (AT5G23575).

Another gene found in the 10-kb windows of this SNP includes the *SUPPRESSOR OF GENE SILENCING 3 (SGS3)* gene (Fig 3B).

**Fig 3.**
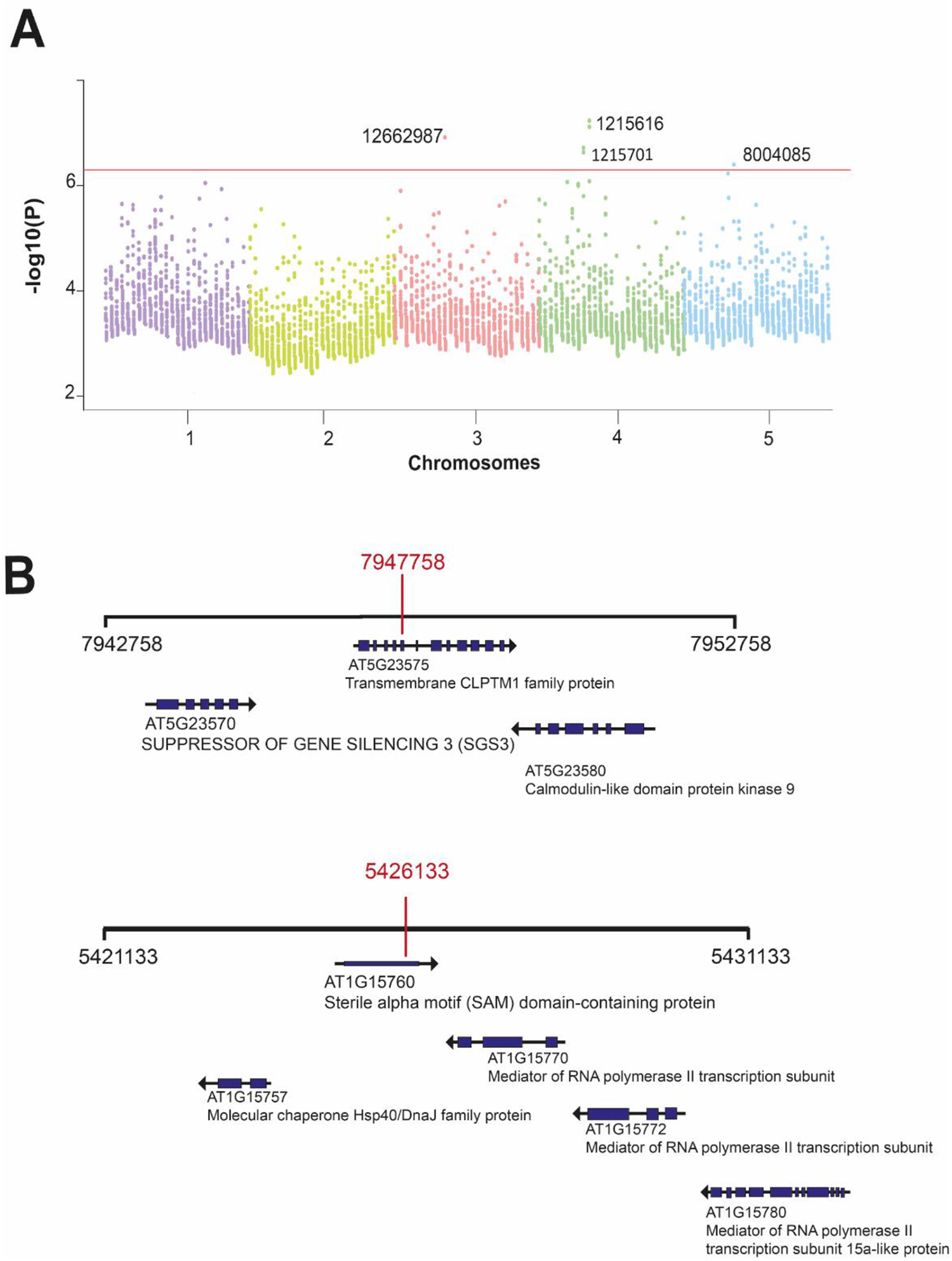
SNPs obtained from individual GWAS and SNPs associated to the genes involved in PR length. (A). Manhattan plot using the top 0.1% SNPs (n=205 SNPs for each study, 5,535 SNPs in total) and their association [-log10(P-value)]. The 10% false discovery rate (FDR) threshold after the Benjamini-Hochberg correction is plotted as a red horizontal line. (B). 10-kbp region surrounding SNPs whose associated genes show significant changes in PR length.

### Genes identified by the distinct GWAS strategies

The top significant SNPs identified by GWAS are commonly not the causal SNP even when it is positioned into gen but rather indicate the genomic region in which the causal gene is located [22], so we select the top SNPs obtained and shared by the data normalized by z-score, case-control, single GWAS, and the pairwise analysis and screened genes within the 10 kb region comprising the associated SNPs detecting forty-one genes in this window (Table 1). Six of these genes have already been shown to be involved in PR growth (Table 3). Near the SNP 8004085, we located the *EARLY FLOWERING BY OVEREXPRESSION 1 (EFO2)* gene that encodes a Transducin/WD40 repeat-like superfamily protein [32]. Although the primary function attributed to this gene has been related to flowering, the *efo2-1* mutant has a shorter PR compared to WT [32]. In this genomic region, we find the *PROPYZAMIDE-HYPERSENSITIVE 1 (PHS1*) gene that encodes a protein similar to the mitogen-activated protein kinase (MAPK) phosphatases. The *phs1-1* is a semi-dominant allele that disrupts normal microtubule functions and has left-handed helical growth and shorter PR than WT. In the homozygous state, the T-DNA insertion in *PHS12* (*phs1-2*) leads to a recessive lethal phenotype, whereas the *phs1-2* heterozygous plants are similar to WT [33,34].

In the SNP 11435245, we detected the gene AT1G31870, a homolog of yeast Bud13p that participates in pre-mRNA splicing [35]. It has been reported that the transgenic lines using an artificial microRNA that targets *AtBUD13* have less transcript accumulation and showed defective seed development and shorter PR in 7-day-old seedlings [35].

One interesting gene that does not have shorter PR compared to WT under control conditions, but it does under ABA treatment, is *SUPPRESSOR OF GENE SILENCING 3 (SGS3)* [36]. SGS3 is essential for producing small interfering RNA of the trans-acting siRNA pathway, which is required to mediate the auxin response in the meristematic zone [37]. This gene was identified in the analysis with the correlation between D5 and D6 (SNP 7947758).

The SNP 18054594 mapped near the *MULTIUBIQUITIN CHAIN BINDING PROTEIN* or *RPN10*, a non-ATPase subunit, part of the 26S proteasome [38]. The T-DNA insertion mutants *rpn10-1* and *rpn10-2* in a C24 and Col-0 background, respectively, have pleiotropic defects as reduced shorter PR length, less inhibition by kinetin and auxin and more sensitivity to ABA [38,39]. The positive correlation finding between the SNP identified by GWAS and the size root phenotype of the overexpression or loss of function mutants in genes associated with these SNPs, support our results (Table 3).

For the genes obtained whose function has not been proved experimentally, we investigated their expression profile in PR using the BAR eFP browser. According to the level of expression in the PR and the available information of their mutants, we selected ten of the forty-one genes found in the different GWAS strategies for further studies: *VP2* (AT1G78920), F-box/RNI-like superfamily protein (AT4G02740), *EFO2* (AT5G23730), AT1G31860 *HISTIDINE BIOSYNTHESIS 2* (AT1G31860), *MBP1* (AT4G38630), *SGS3* (AT5G23570), *CDPK9* (AT5G23580), SAM domain-containing protein (AT1G15760), AAR2 protein family (AT1G66510) and *NRPD2A* (AT3G23780). From these, only the loss of function mutants from EFO2 and MBP1 display shorter PR than WT under control conditions (Table 3). The spatial expression of the ten candidate genes is presented in S2 Fig. All the genes were expressed in the PR, mainly in the meristematic zone. For the SAM domain-containing protein and the F-box/RNI-like superfamily protein, the only available information was obtained from Klepikova Atlas, showing high expression in the radicle (S2 Fig).

**Table 3.** Genes described in PR development and their phenotype in overexpressing (OE) or loss of function (LoF) mutants.

| Gen | Phenotype | Function | Reference | Score obtained in our analyses |
| --- | --- | --- | --- | --- |
| AT1G75590 *SMALL AUXIN-UP 52 (SAUR 52)* | Longer PR in OE mutants | Member of the Primary auxin-responsive (PAR) genes, rapidly induced in response to auxin | [40] | 5.7 |
| AT5G23730 *EARLY FLOWERING BY OVEREXPRESSION 2 (EFO2)* | Shorter PR in LoF mutant *efo2-1* | Repressor in photoperiodic flowering. The LoF mutant shows vegetative growth perturbations | [32] | 6.39 |
| AT5G23720 *PROPYZAMIDE-HYPERSENSITIVE 1 (PHS1)* | Shorter PR and left-handed helical growth in *phs1-1*, a semi-dominant allele | A phosphatase that regulates MAPK and is involved in the organisation of cortical microtubules. | [33,34] | 6.39 |
| AT1G31870 Arabidopsis homolog of yeast Bud13p (Bud site selection protein 13) AtBUD13. | Shorter PR in a line where an artificial microRNA targets AtBUD13 | Participation in pre-mRNA splicing | [35] | 5.2 |
| AT5G23570 *SUPPRESSOR OF GENE SILENCING 3 (SGS3)* | Shorter PR in *sgs3* LoF mutant under ABA treatment | Protein is necessary for the production of small interfering RNA of the trans-acting siRNA pathway. It is required to mediate the auxin response in the meristematic zone | [36] | 5.77 |
| AT4G38630 *MULTIUBQUITIN CHAIN BINDING PROTEIN (MBP1/RPN10)* | Shorter PR in the *rpn10-1* and *rpn10-2* LoF mutants | A non-ATPase subunit, part of the 26S proteasome | [38,39] | 6.06 |

### NOJO and SGS3 genes discovered via GWAS regulate root length

To investigate the participation of these 10 genes in the PR length, we used T-DNA lines ordered from NASC. The genotypification was performed using a left border primer of the T-DNA insertion and primers flanking the insertion (S10 Table). We only obtained homozygous lines for *AVP2*, F-box/RNI-like superfamily protein, *CDPK9*, SAM domain-containing protein and *SGS3.* For *HISN2*, we used the heterozygous line to evaluate the PR length. *HISN2* encodes a protein with two domains that function as a cyclohydrolase and a pyrophosphohydrolase and participate in histidine biosynthesis; knockout alleles of this gene are embryo-lethal (Muralla et al., 2007). For that reason, we genotypificated 90 plantlets of *hisn2* and associated the PR length at 12 DAS with the genotype (heterozygous or WT). We divided the PR length into short, medium and long PR according to Sturge’s rule to see the distribution of the values (short PR length: 3.949-5.727, medium PR long 5.728-7.505 and PR length:7.506 9.283 cm (S3 Fig). We did not find a clear association between the presence of the insertion and the PR length; from seedlings with large PR only 61% have the insertion compared to the 38% which do not have it. Further studies are required to validate if *HISN2* participates in PR growth.

For the homozygous lines, we carried out PR length kinetics over 12 days, starting from day two after sowing. From these, the T-DNA line SALKseq_056622.2, which interrupts the single exon of the gene that encodes a SAM domain-containing protein and *sgs3-13 (*SALK_039005) show a larger PR length than WT (S4 Fig). To confirm these data, we order a second allele for these genes, SALK_208945C, for SAM domain-containing protein, whose T-DNA line is inserted in the first exon and for *SGS3*, we used the *sgs3-11* line that has a splice site mutated from G to A [41]. For both alleles we performed kinetic growth for 12 days and we observed that the PR length is larger than WT, confirming that these genes are negative regulators of PR growth (Figs 4A and B, S5-6 Figs). To verify the expression levels of these genes, we extracted RNA from the T-DNA lines and Col-0 and evaluated by qPCR the transcript levels of 12 DAS roots. We observed reduced expression levels for both alleles of the SAM domain-containing protein and *sgs3-13* compared to WT. For *sgs3-11* we only checked the reported phenotype for this mutant with the loss of leaf polarity [41]. Because the gene that encodes the SAM domain-containing protein has not been described, we named it *NOJOCH MOOTS (NOJO)* which in Maya, means big root.

**Fig 4.**
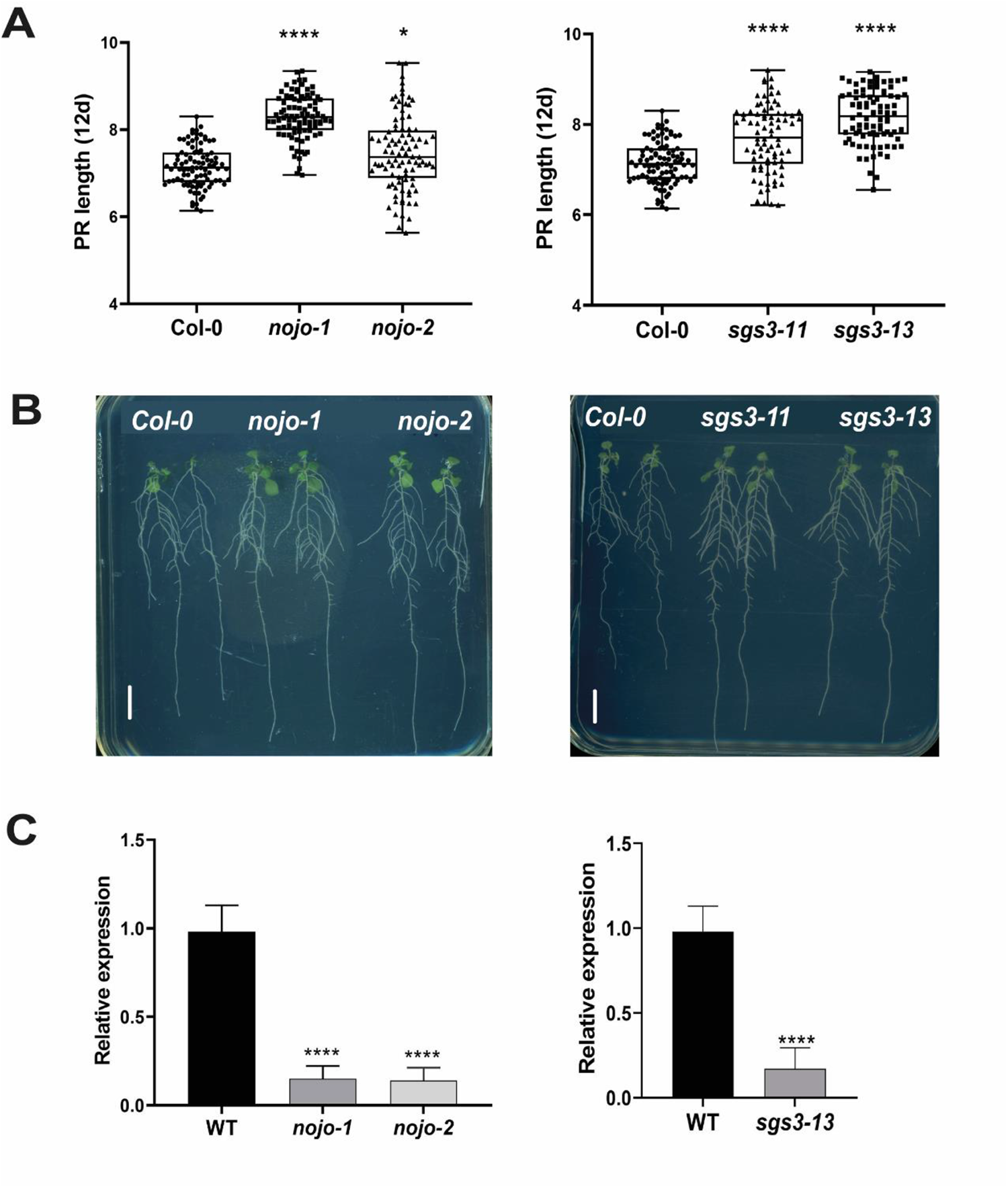
*NOJO* and *SGS3* are novel genes involved in PR length. (A) PR length at 12 DAS of two alleles of Col-0, *SGS3* and *NOJO* (*n* = 90 seedlings used per allele). Significant value is indicated (***) *p* < 0.001, (*) p< 0.05 using ANOVA one-way followed by Tukey’s post-hoc tests (B) Representative seedling at 12 DAS of two alleles of *NOJO* and *SGS3* that were grown in the same plate with Col-0. Scale bar = 1 cm. (C) Relative expression of SAM domain-containing protein (*NOJO)* and *SGS3* in two T-DNA lines. Data represent the means ± SD (n =9). For *nojo* lines and WT, ANOVA one-way followed by Tukey’s post-hoc tests were performed, for *sgs3-13* and WT, unpaired two-tail Student t test was used (***) *p* < 0.001.).

### Polymorphism patterns around the SGS3 and NOJO in extreme accessions

To further analyzed *SGS3* and *NOJO* genes, we searched for the SNPs using representative extreme accessions employing 15% of accessions with short root phenotype and 15% of accessions with long root phenotype that were shared for more than two studies (S8 Table) and their sequences were available in the 1001 Genomes Project.

In the case of SGS3, we detected that 62.5 % of the accessions with short PR lengths display several SNPs in their first and second exon and the first intron compared to only 7.1% of the accessions with long PR lengths. These changes are classified in Table 4; one of these changes is a missense variant at the second exon (CDS position: 1069) in the XS (named after rice gene X and SGS3) domain which is a single-stranded RNA binding domain [42]. This variant produces an amino acid change (M/L, both are non-polar aliphatic amino acids) in 35% of the accessions analyzed with a short PR (ROU-0, HR-5, NFA-8, NFA-10 and SQ-1). In contrast, none of the accessions with long PR length have these SNPs (Fig 5, Table 4). Another SNP in 35% of accessions short PR length occurs at the boundary of an exon and an intron leading to a splice acceptor variant. This kind of change can disrupt RNA splicing resulting in the loss of exons or inclusion of introns and, consequently, an altered protein-coding sequence [43] (Table 4).

**Fig 5.**
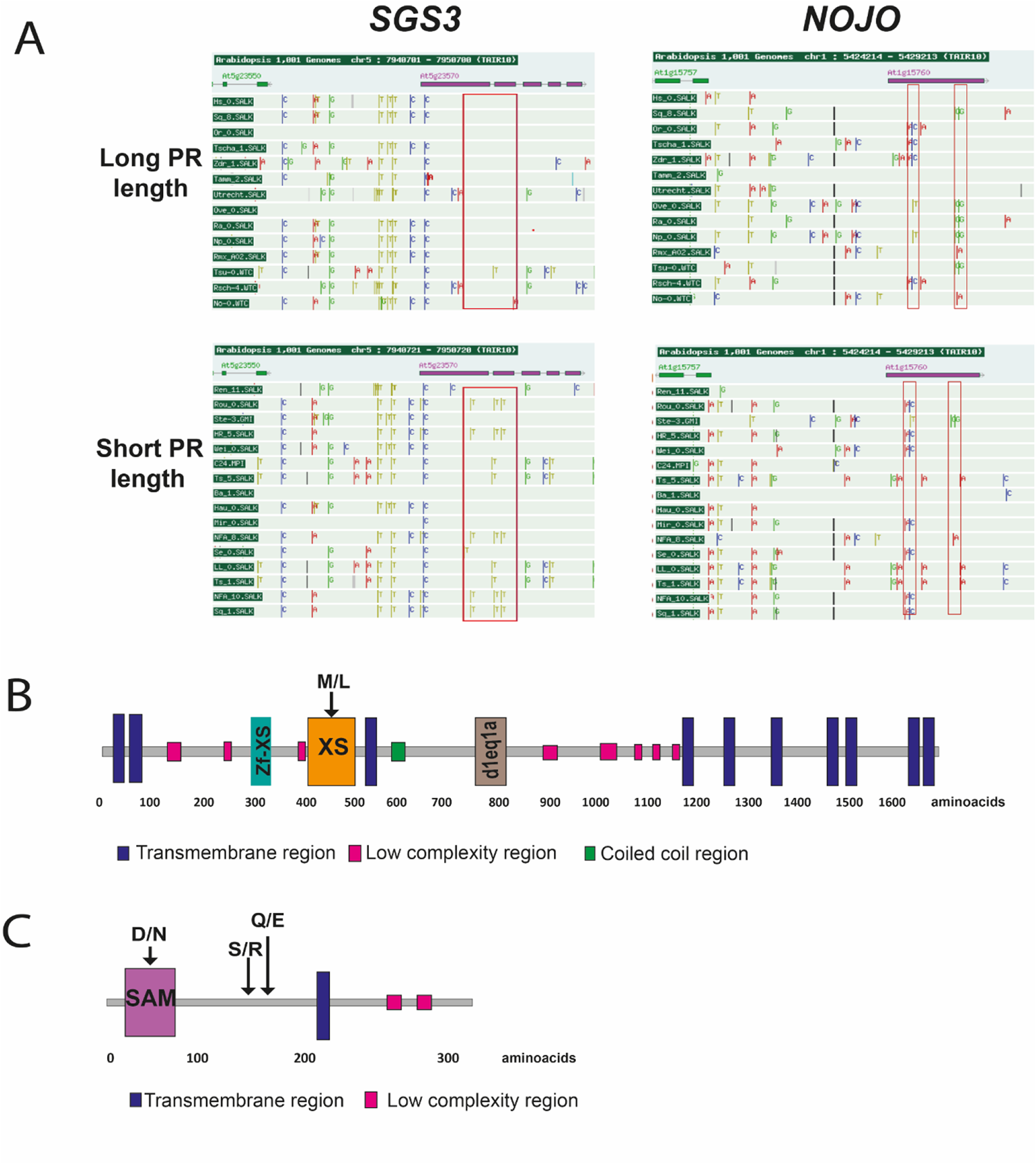
SNPs observed in *SGS3* and *NOJO* from accessions with large and short PR length. **(A)** The red rectangles indicate the SNPs found in *SGS3* and *NOJO* genes and each nucleotide displays different colours. In *SGS3*, the main changes are kept in the first and second exon and first intron in accessions with a short PR. In the gene that encodes a SAM domain-containing protein, the SNPs are detected in the distal part of the single exon in some accessions with long PR length and in the proximal part of the exon for accessions with short PR. (B) SGS3 protein. The amino acid change is indicated in the XS domain. (C) NOJO protein. The amino acid changes were found in the SAM domain and in a sequence without a described domain. In B and C, the main domains are depicted with different colours, and regions with uncommon domains are shown in grey.

For *NOJO,* we identified two missense variants in its exon (CDS position 420 and 445) in 35% of accessions with large PR length (SQ-8, OVE-0, RA-0, NO-0 and TSU-0), whereas only one of the accessions with short PR (STE-3) has this polymorphism. Another missense mutation is found at the SAM domain (CDS 118) in 43.7% of accessions with short PR compared to 28.5% of accessions with long PR, and this SNP led to a D/N change (acidic amino acid to a polar aliphatic amino acid) (Table 4, Fig 5C). The SAM domain functions in protein-protein interaction and it is common in scaffolding proteins, transcription regulators, tyrosine kinases and serine/threonine kinases [44].

**Table 4.** SNPs found in contrasting accessions.

| *Gene* | *Position* | *Nucleotide change* | *Amino acid change* | *PR length* | *Percentage of accessions* |
| --- | --- | --- | --- | --- | --- |
| AT1G15760<br>*NOJO* | CDS 118<br>First exon. SAM domain | Missense variant<br>**GAC/AAC** | D/N | SHORT | 43.7% |
|  |  |  |  | LARGE | 28.5% |
|  | CDS 150<br>First exon. SAM domain | Synonymous variant<br>TCT/TCC | S | SHORT | 43.7% |
|  |  |  |  | LARGE | 28.5% |
|  | CDS 420.<br>First exon. | Missense variant<br>**AGC/AGG** | S/R | SHORT | 6% |
|  |  |  |  | LARGE | 35 % |
|  | CDS 445.<br>First exon | Missense variant<br>**CAA/GAA** | Q/E | SHORT | 6 % |
|  |  |  |  | LARGE | 35 % |
| AT5G23570<br>*SGS3* | CDS 711<br>First exon. zf-XS Domain | Synonymous variant<br>**GCC/GCT** | A | SHORT | 35% |
|  |  |  |  | LARGE | 0% |
|  | First intron<br>5:7944656 | Splice acceptor variant C/T | - | SHORT | 35% |
|  |  |  |  | LARGE | 0% |
|  | CDS 1069<br>Second exon. XS Domain | Missense variant<br>**ATG/TTG** | M/L | SHORT | 35% |
|  |  |  |  | LARGE | 0% |
|  | First intron<br>5: 7944632 | Intron variant | - | SHORT | 7% |
|  |  |  |  | LARGE | 25% |

## Discussion

Arabidopsis has been used as a model to understand the genetic bases of root growth due to its experimental amenability. In many studies, seedlings are grown on sterile semi-solid agar media in Petri dishes allowing non-invasive observations. However, the conditions can vary between research groups, and Arabidopsis plants can be cultivated in unnatural conditions. Only shoots are illuminated in nature, whereas roots grow in darkness; however, under standard agar-plate culture, both are illuminated [45]. We compared the PR length of 639 accessions from nine studies under control conditions but with different growth conditions. Lachowiec et al. (2015) is the only study that grows the accessions in darkness and shows the highest mean PR length [23]. In contrast, the majority of the studies used a 16/8-h light/dark cycle [11,13,14,16–18] or 12/12 h light/dark [12]. It has been reported that the quality, quantity, type and direction of light affect the PR length [46].

Besides, it has been published that sucrose addition may cause artefact root responses [45] and almost all the studies that we analyzed added about 0.5% – 2% sucrose into the agar medium, except Lachowiec et al. (2015) that use a sucrose-free medium [23]. Additionally, we noted that the Murashige and Skoog (MS) medium differs between studies, ranging from 0.2 to 1X, and it has been described that the concentration of the MS medium influences the growth of the radicle: the radicle length rises as the MS concentration increases and then declines afterward [47]. Thus, the impact of the medium composition on PR length is particularly relevant since it depends on nutrient availability [48]. Recently, it was proposed an optimized *in vitro* protocol that uses an optimal nutrient balance in the medium close to natural soil conditions to avoid biases in phenotypic observation [49]. A good example of this phenotypic variation due to growth conditions is the Kin-0 accession, the only one that is shared between the nine studies we found and displays ample PR length even if the same day is evaluated. For that reason, we remarked that the length of the PR must be indicated in the different studies and not only the DAS or DAG.

We pointed out that the data of Deolu-Ajayi, (2019) does not correlate with any other studies due to the conditions used, such as the photoperiod (12h/h), MS 0.5 and 0.5% sucrose [12]. In their study, they showed the light intensity (122 µmolm-2s-1), and although it is the standard, in other articles, the light intensity is not specified. These conditions render a faster growth rate compared to the other studies that carry out growth kinetics, still when we examine the same accessions.

We have shown that the experimental conditions are crucial to determine PR length as we have found contrasting phenotypes for the same accession under the different studies. This might sound evident, but we have not found any manuscript that demonstrated this before. Besides, the accessions used are different between studies making a comparison of PR length, almost impossible. Therefore, we performed different GWAS tests to check for SNPs that could provide novel genes that participate in PR length as follows: 1) using the data of single studies, 2) making pairwise analysis with high correlation associations, 3) analyzing the same number of accessions in different studies and 4) using the daily data of the root growth kinetic of the same research. We all of these methodologies, we found 41 candidate genomic loci (Table 3), of which six have been described as promoters or repressors of PR growth [32–35,38–40].

According to our results, the best strategy to retrieve causal SNPs was obtained in the pairwise analysis comparing the DAG or DAS with higher correlation in the same study and selecting the common SNPs highly scored [16,24,50]. Moreover, it has been reported that during PR development, the score of the same SNPs obtained by GWAS change emphasising the need to perform PR course growth. Accordingly to what has been shown for Arabidopsis root architecture, we found that the associations with some SNPs increase while others decrease during PR growth [16,24,50]. In addition, the same accessions can be compared when they have a high correlation even though the seedlings have varying ages, and recover reliable candidate SNPs (S9 Table). Contrary, when we compared the same accessions, but with a low correlation, we did not recover significant SNPs.

In this study, we found two novel genes that participate in PR length: a gene that encodes a Sterile Alpha Motif (SAM) domain-containing protein (AT1G15760) that has not been described before but with a high expression in PR, which we named *NOJO*, and *SGS3,* that is involved in RNA-mediated gene silencing and it is also express in PR.

In Arabidopsis, 12 SAM domain-containing proteins have been annotated, and the best-characterised protein with this domain is LEAFY (Denay et al., 2017). According to the sequence-based prediction, NOJO displays a subcellular localization in chloroplast and can interact with several proteins predicted to be involved in vesicle trafficking processes, including SNARE-associated proteins and vesicle transport family proteins, suggesting its function in endomembrane trafficking [51]. In this study, we are showing a function of one of these SAM proteins as two knockdown alleles of *NOJO* showed longer PR.

On the other hand, *SGS3* a well-studied gene that participates in the biogenesis of trans-acting small interfering RNAS (ta-siRNAs), is involved in leaf polarity [52]; lateral root initiation [53] and during the response modulation of the transit-amplifying region of the root meristem to exogenous auxin [37]. Contrary to what has been reported that *SGS3* does not participate in primary root length under control conditions [36,54] we found that *sgs3-11* and *sgs3-13* show a longer PR at 12 DAS, indicating SGS3 participation in primary root length. This could be explained by the developmental stage of the plants as we measured them at 12 DAS. Furthermore, 62.5 % of the accessions with short PR lengths show SNPs in their second and third exon compared to the 7.1% of the accessions with long PR lengths. Besides, 35% of accessions with short PR display a missense variant in the XS domain that is necessary during RNA recognition and it has been reported that although the sequence between the XS domain and other RNA-binding domains is low, the secondary structures is very important as exhibits a high resemblance similarity showing functional conservation. It could be interesting to test if these SNPs change the protein structural fold and have a function during PR growth [42].

The root plastic responses difficult the comparative analyses among data obtained in different growth conditions; however, our approaches using different GWAS tests were successful to find SNP -associate genes involved in root growth, being a method that can be used to obtain genes associated with different traits. Despite the heterogeneous growth conditions, we find two novel genes that are repressors of primary root growth.

## ACKNOWLEDGMENTS

We sincerely thank Dr Marcos Castellanos for providing us with the T-DNA lines, Elsa H Quezada-Rodríguez for helping us with the protein analysis and Dra. Diana B. Sanchez Rodríguez for her logistical and technical support.

## Supporting information

S1 Fig. Histograms of the published data

S2 Fig. Root expression of candidate genes found in GWAS

S3 Fig. PR length division of *his2* heterozygous line

S4 Fig. PR length kinetics over 12 days of homozygous T-DNA lines from candidate genes obtained from GWAS

S5 Fig. PR length kinetics over 2-12 days of *nojo-1, nojo-2, sgs3-11* and *sgs3-13*.

S6 Fig. Growth rate over 2-12 days of *nojo-1, nojo-2, sgs3-11* and *sgs3-13*

S1 Table. Growth conditions and accessions used in each study

S2 Table. Primary root length from all studies

S3 Table. Maximum and minimum values from PR length in all accessions.

S4 Table. Frequency of accessions used in each study

S5 Table. Normality test

S6 Table SNPs obtained in GWAS using data normalised by Z-score

S7 Table. SNPs shared between studies found in individual GWAS of all the studies

S8 Table. 15% of accessions with shorter and longer PR from each study.

S9 Table. Pairwise GWAS in inter and intra-studies with high correlation

S10 Table. The T-DNA insertion lines of the candidate genes and primers used

## Author Contributions

BAL-R, AG-A and AU-A conceived and wrote the study. BG-P, MPS and EA-B helped in writing and editing the article. All authors have read and agreed to the published version of the manuscript.

## Funding statement

This study was financed with the following grants: CONACyT: 102959 and 102987 (MPS) and AG-A, BG-P, MPS and EA-B received funding from UNAM-DGAPA-PAPIIT (IN200920, IN206223, IN203223, IN211721). BAL-R received a postdoctoral fellowship from CONACyT (489444).

## Notes

https://doi.org/10.5281/zenodo.8156775

### Competing Interest Statement

The authors have declared no competing interest.

